# Mathematical reconstruction of the metabolic network in an *in-vitro* multiple myeloma model

**DOI:** 10.1101/2022.09.12.507672

**Authors:** Elias Vera-Siguenza, Cristina Escribano-Gonzalez, Irene Serrano-Gonzalo, Kattri-Liis Eskla, Fabian Spill, Daniel Tennant

## Abstract

It is increasingly apparent that cancer cells, in addition to remodelling their metabolism to survive and proliferate, adapt and manipulate the metabolism of other cells. This property may be a telling sign that pre-clinical tumour metabolism studies that exclusively utilise *in-vitro* mono-culture models could prove to be limited for uncovering novel metabolic targets that can translate into clinical therapies. Although this is increasingly recognised, and work addressing this is becoming routinary in a rapidly emerging field, much remains unknown.

This study employs an interdisciplinary approach that leverages the predictive power of mathematical modelling to enrich experimental findings. We develop a functional multicellular *in-silico* model that facilitates the qualitative and quantitative analysis of the metabolic network spawned by an *in-vitro* co-culture model of bone marrow mesenchymal stem- and myeloma cell lines. To procure this model, we devised a bespoke human genome constraint-based reconstruction workflow that combines aspects from the legacy mCADRE & Metabotools algorithms, the novel redHuman algorithm, along with ^13^C-metabolic flux analysis. Our workflow transforms the latest human metabolic network matrix (Recon3D) into two cell-specific models coupled with a metabolic network spanning a shared growth medium. When cross-validating our *in-silico* model against the in-vitro model, we found that the *in-silico* model successfully reproduces vital metabolic behaviours of its *in-vitro* counterpart; results include cell growth predictions, respiration rates, as well as support for observations which suggest cross-shuttling of redox-active metabolites between cells. Together, our methodology and its results provide yet another step toward the relevance of studies of this type in the field.

## Introduction

A cell’s metabolism can be defined by extensive interconnected chemical reaction networks subdivided by seemingly organised pathways [1, 2]. These pathways specialise in building unique blocks necessary for all aspects of cellular functioning (i.e., homeostasis, cell reproduction, and biomass) [3, 4]. Their performance depends on many aspects, a critical one being the microenvironment in which the cell resides [4–7]. A microenvironment consists of a reservoir of readily available chemicals and different cell types. As the microenvironment’s cell population have different metabolic requirements, their spatial organisation, type, and objectives will be affected by the properties of the microenvironment itself. This is particularly evident in the case of cancer metabolism [8]. Studies have demonstrated that several types of malignancies rely on their immediate extracellular environment and the cells that reside within to meet their highly energetic demands, enhance proliferation, and even enable therapy resistance [5–7, 9–13].

A powerful tool for probing the metabolic interactions between cell types are *in-vitro* co-cultures (Fig. 1) [10–13]. These experimental models are advantageous as cell interactions can be studied under a controlled microenvironment that allows the co-cultured cells to establish a unique metabolic niche [14–16]. The resulting intercellular chemical exchanges enable cells in the co-culture to tap into a larger pool of metabolites and, thus, instigate a substantial division of metabolic labour that aims to optimise the use of available nutrients. Consequently, cell survival and proliferation may be enhanced, particularly under system perturbations (e.g., hypoxia or lack of oxygen) in which mono-cultures would otherwise perish [14–16].

**Fig 1.**
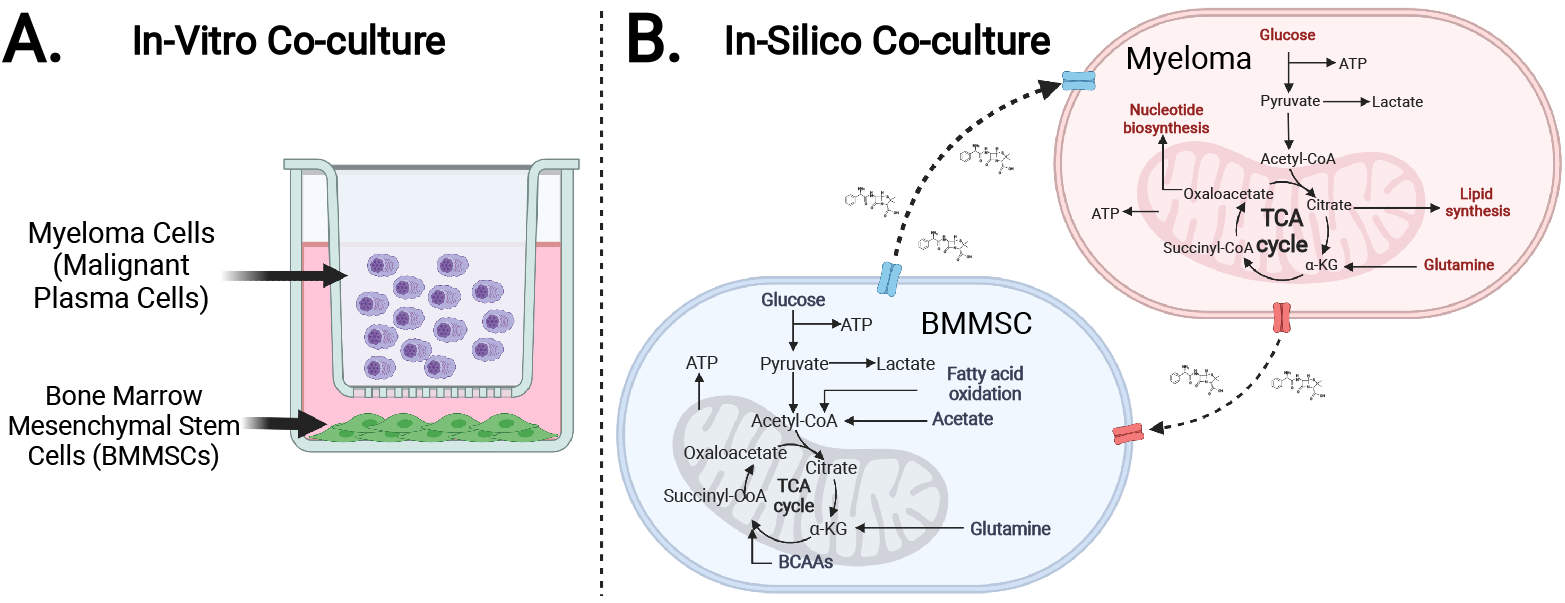
Schematic diagram depicting the *in-vitro* co-culture model setup. (*Created with BioRender*.*com*)

Co-culture studies are possible today thanks to the technological advancement seen during the past couple of decades [17]. Nevertheless, the study of intercellular metabolic interactions across microenvironments remains incredibly challenging [18–20]. As a result, a wave of interdisciplinary research avenues has given rise to numerous bioinformatic approaches that unravel cell metabolism. Techniques like omic enrichment and extensive multi-scale and multi-omics analyses have helped underpin critical aspects of cell functionality with unprecedented precision [21–23]. However, the granularity required to fully understand the complexities of cell biochemistry, such as quantitative information on metabolic fluxes, is, more than often, too challenging to obtain, analyse, and interpret.

This is an all-too-familiar problem in multiple myeloma (MM) research. MM is a type of malignancy found in the bone marrow (BM) and is formed through the ingress and proliferation of malignant plasma cells [11, 24]. The disease remains incurable, and even though novel therapies have improved patient survival outcomes, more effective treatments are needed for many patients unable to tolerate current cancer management strategies [25, 26]. Therefore, developing next-generation therapies for multiple myeloma requires a technical approach focusing on delivering long-term disease control while maintaining the quality of life [26].

A potential approach to address this issue involves exploiting those metabolic features critical for MM survival. Malignant plasma cells have been observed to transform the phenotype of the stroma residing in the bone marrow niche [7, 15, 27]. This malignancy can generate a metabolic network within the tumour microenvironment capable of promoting survival, and aggressive proliferation [28, 29]. Studies have previously demonstrated that the metabolism of the bone marrow is significantly altered in patients with MM and that the bone marrow mesenchymal stem cell (BMMSC) is a primary supportive cell type for malignant plasma cells [28, 30]. We thus hypothesise that understanding the physiology of the metabolic interaction between these two cell types (the BMMSC and the MM cell) and the niche in which they reside will facilitate the discovery of a novel metabolic target. Such a target could interfere with a metabolic intercellular pro-survival axis presumed to be critical for the malignancy’s survival.

In this light, our study outlines a multidisciplinary and multifaceted approach to produce a testable and integrated *in-vitro/in-silico* model of the metabolic network formed by malignant plasma cells, or myeloma, and bone marrow mesenchymal stem cells. We achieve this via two phases that display the synergy between computational and bench-side experimental research. The first or experimental phase involves mono-culturing HS-5 (BMMSC) and JJN-3 (MM) cell lines. We performed benchmark measurements such as cell growth and cell respiration rates, that is, oxygen (O_2_) consumption, to probe their carbon metabolism capacity [31, 32]. We followed these experiments by *in-vitro* co-culturing the aforementioned cell lines (Fig.1 A). Finally, we probed the metabolic phenotype of the co-culture system using stable-isotope ^13^C_6_Glucose and ^13^C_5_Glutamine tracing [33, 34].

The computational phase of this study draws from and optimises established techniques to generate individual cell-specific genome-scale models (GEMs) of our two cell types, the BMMSCs, and the MM [35, 36]. Using these GEMs together, we reconstruct an *in-silico* co-culture constraint-based model (CBM). The model is assembled through a bespoke computational workflow that integrates publicly available transcriptomic data sets (retrieved from the NCBI Gene Expression Omnibus - GEO), using the mCADRE algorithm, to a thermodynamically constrained version of the latest global knowledge base of metabolic functions categorised for the human genome: the Human Recon3D; using a subroutine of the redHuman algorithm (Fig. 2) [37–40]. Our computational co-culture was further constrained to mimic the *in-vitro* growth medium, RPMI, using the metabotools MATLAB routine. We validate the resulting model against its experimental counterpart, where we found it successfully recapitulated the observed *in-vitro* growth and respiration rates. Furthermore, via integration of stable-isotope labelling data via ^13^C-metabolic flux analysis (^13^C-MFA), using the MATLAB INCA routine as a means of “phenotype tuning” [41, 42].

**Fig 2.**
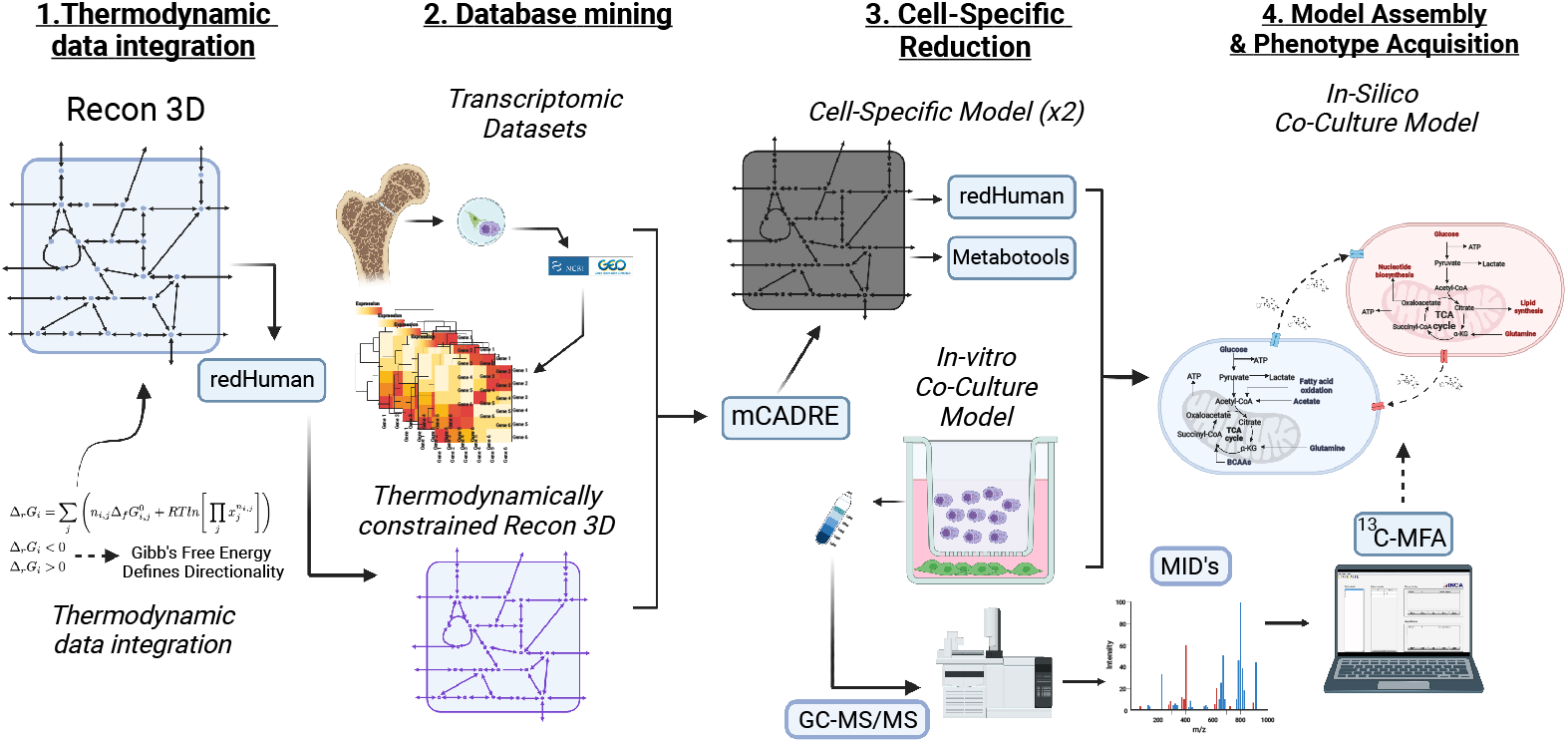
Schematic diagram depicting the workflow to generate our *in-silico* co-culture model.

## Materials and methods

### Cell culture

The HS-5 cell line (ATCC) and JJN-3 cell lines were both STR profiled to ensure accurate identification and maintained in RPMI (Sigma-Aldrich, R8758) with 10% FBS (Sigma-Aldrich, F7524). For individual cell growth experiments, HS-5 and JJN-3 cells were seeded at 4×10^4^ cells/mL, and 3.7×10^4^ cells/mL and cell growth was assessed over time by cell count for JJN-3 and by sulforhodamine B (SRB) assay for HS-5. For co-culture experiments, HS-5 cells were seeded at 12×10^4^ cells per well of a 6-well plate the day before JJN-3 were seeded on top of a transwell (CC401, Appleton Woods) at 8 × 10^4^ cells per well.

### Oxygen consumption

Trypsinized cells were resuspended in culture media and loaded in an Oxygraph-2k (Oroboros instruments) chamber. After closing the chambers and recording routine respiration, Oligomycin (2.5 *µ*M) was added to inhibit ATP synthase. Respiration was inhibited by the addition of Rotenone (0.5 *µ*M) and Antimycin A (2.5 *µ*M) at the end of the experiment. Measurements of the non-phosphorylating electron transfer system (ETS) capacity were obtained through stepwise (0.5 *µ*M) titration of the uncoupler, carbonyl cyanide 4-(trifluoromethoxy) phenylhydrazone (FCCP).

### ^13^Carbon metabolic tracing

Cells at 70% confluency were incubated in flux media: modified RPMI 1640 without glucose, glutamine, L-isoleucine, L-leucine, L-valine, and phenol red (Cell Culture Technologies) supplemented with either 2g/L ^13^C-[U]-glucose or 2mM ^13^C-[U]-glutamine (both CK isotopes), with the other provided in an unlabelled form. After 48 hours, media was removed for extraction, JJN-3 cells spun down, and then cells (JJN-3) and wells (HS-5) were washed twice with ice-cold saline before quenching metabolism with ice-cold MeOH. Cells were transferred to a cold tube into which was added D^6^-glutaric acid in ice-cold H_2_O (CDN isotopes, D-5227) and pre-chilled chloroform. After shaking on ice for 15 minutes and centrifugation, the polar phase was transferred to another tube to be dried.

### Derivatisation and GC-MS

Dried-down extracts were derivatized with 2% methoxamine in pyridine (20*µ*l, for one hour at 60°C) followed by N-(tert-butyldimethylsilyl)-N-methyl-trifluoroacetamide, with 1% tert -butyldimethylchlorosilane (30*µ*l, 1h at 60°C). Samples were transferred to glass vials for GC-MS analysis using an Agilent 8890 GC and 5977B MSD system. 1*µ*L of the sample was injected in splitless mode with helium carrier gas at a 1.0 mL/min rate. The initial GC oven temperature was held at 100°C for one minute before ramping to 160°C at a rate of 10°C/min, followed by a ramp to 200°C at a rate of 5°C/min and a final ramp to 320°C at a rate of 10°C/min with a five-minute hold. Compound detection was carried out in scan mode. Total ion counts of each metabolite were normalised to the internal standard D^6^-Glutaric acid.

### Normalisation and quantification

Data normalisation occurred by counting cells using counting chambers Fast Read 102, purchased from Kova International. GCMS data were analysed using Agilent Mass Hunter software for real-time data quality analyses and in-house MATLAB scripts.

## Model

A frequently sought-after solution for producing *in-silico* large metabolic models is to map out tissue-specific functions using genome-scale models (GEMs) [43, 44]. GEMs are comprehensive models that aim to recreate a particular organism’s metabolism [36, 38, 43, 44]. They typically include most, if not all, known chemical reactions and metabolites in the form of a stoichiometric matrix and their corresponding genes in the form of a gene-rule matrix. The latter is an attractive feature of GEMs because each enzyme-mediated reaction in the model is associated with information regarding how a gene influences a protein, which in turn influences a reaction. In GEMs, this is referred to as a gene-protein-reaction (GPR) rule. GPRs are essential to GEMs as their description of the relationship between genes encoding enzymes enables the mediation of a given reaction or set of reactions in a given model [43, 45]. Furthermore, GEMs are attractive because they promise to advance our understanding of the underlying metabolism behind various physiological and pathological processes in various tissues [43, 45, 46]. Unlike many modelling frameworks, GEMs can yield a profound insight regarding cell behaviour requiring minimal information on the biophysical equations that require difficult-to-measure kinetic parameters. Efforts such as the global knowledgebase of metabolic functions categorised for the human genome, known as Human Recon3D, coupled with abundant high-throughput data, make the reconstruction of cell-specific metabolic models possible [39, 47].

What follows is a detailed workflow of a top-down reconstruction strategy designed to produce two cell-specific constraint-based models; one bone marrow mesenchymal stem cell (BMMSC) and one myeloma (malignant plasma - MM) cell (Fig. 2). Our overarching aim is to assess the consequences of metabolic adaptation and regulation of the desired phenotype - in our case, a multiple myeloma *in-vitro* co-culture; via network flux analysis. To this end, the first step uses the generic model of human metabolism: Recon3D, the latest and arguably the most comprehensive consensus human GEM. Recon3D consists of 10,600 reactions, 5938 associated with 2248 genes and 2797 unique metabolites across seven compartments: cytosol, mitochondria, peroxisome, Golgi apparatus, endoplasmic reticulum, nucleus, and lysosome [39, 47]. GEMs of metabolic networks combined with CBM produce a powerful tool that ultimately yields qualitative and quantitative information regarding the distribution of steady-state metabolic fluxes through a given network graph representation. This feature enables the prediction of experimentally measurable variables such as cell growth rates, or production rates of metabolites of interest [48]. In this paradigm, the steady-state approach works because, within the time scale of our experiments, changes in the concentration of metabolites occur slowly [48–50]. This is even true under constant exponential cell growth (i.e., cancer). The steady-state assumption allows CBMs to model large networks without knowing enzyme kinetics and post-translational regulatory mechanisms [49]. This last feature is advantageous when cellular activity in interest physiology is not well-understood [49, 51]. In the appendix section (A. 1) of this article, we give a brief overview of the mathematical formulation of CBMs and flux analysis.

### Thermodynamic data integration (Step 1)

We introduced thermodynamic information for all metabolite compounds and reactions by following the protocol outlined in the redHuman workflow [40]. Briefly, the protocol involves obtaining metabolite Gibb’s free energy data from the MetaNetX archive [52]. Those metabolites from Recon3D with identifiers from SEED, KEGG, CHEBI, and HMDB, are manually annotated and then, using ChemAxon’s Marvin (available for free if used for academic purposes), the compound structures were transformed into their primary protonation states at a pH of 7 and generated MDL Molfiles [52–58]. These MDL molfiles and the Group Contribution Method were then used to estimate the standard Gibb’s free energy of the formation of the metabolites in Recon3D as per [59, 60]. As a result, each metabolite is associated with its Gibb’s free energy (Δ*G*). Then the Δ*G* of a given reaction is estimated from the thermodynamic properties of its reactants and products. Following the incorporation of this data, the model was then primed for thermodynamic flux balance analysis (tFBA) and thermodynamic flux variability analysis (tFVA) using the prepModelforTFA.m routine from redHuman [40]. A readily available annotated matrix and a MATLAB script are available in the matTFA package by [61]. We, however, repeated the protocol due to updates in the MetaNetX archive (with the latest update occurring in March 2022). The model was verified for consistency by running the redHuman’s matTFA subroutine and comparing its results to the thermodynamically annotated Recon3D version generated by the redHuman algorithm package [40, 61].

Performing this step allows one to determine and ensure the feasible thermodynamical directionality of those chemical reactions included in the Recon3D matrix for which we have no constraining data. Numerically, this also ensures that the employment of optimisation routines, such as optimiseCBmodel from the MATLAB COBRA routine, does not produce thermodynamically infeasible flux loops, thus avoiding physiologically inaccurate behaviour [62].

### Database mining and cell-specificity (Step 2)

To determine which reactions must be removed from the now thermodynamically constrained Recon3D matrix to generate our cell-specific models, we used transcriptomic datasets from the Gene Expression Omnibus database (GEO) [37, 39]. In this study, we adhere to the data sets based on the Affymetrix Human Genome U133 Plus 2.0 Array platform for the bone marrow mesenchymal stem cell model and the malignant plasma cell [63]. The choice was driven by full support from Affymetrix via the availability of the .cdf file (a file that describes the layout for an Affymetrix GeneChip array) and its compatibility with the latest MATLAB version [63]. Those sets for generating the bone marrow mesenchymal stem cell model reconstruction were obtained from healthy human early bone marrow passage samples and gene expression data from mesenchymal stem cells cultured in MSCGM (mesenchymal stem cell growth medium). These were chosen in line with an early model that uses a legacy version of Recon3D, namely Recon1, to generate a genome-scale reconstruction-*iMSC1255* [64, 65]. Their accession ID: GSM184636, GSM184637, GSM184638, GSM194076, GSM194077, GSM194078, GSM194079, GSM764199, GSM797497, GSM797498, GSM920586, GSM920587, and GSE80608 [66]. In particular, the GSE80608 data set includes newer datasets gather from gene expression of bone marrow mesenchymal stem cells grown out from bone marrow aspirates for multiple patient passages [30]. For the myeloma cell model, we have used those datasets from a series of pre-treatment bone marrow aspirates derived from multiple myeloma patients. These consist of exclusively CD138+ expressing plasma cells from monoclonal gammopathy of unknown significance (MGUS) and multiple myeloma bone marrow samples from newly diagnosed and previously untreated patients. Their accession ID: GSM50986, GSM50987, GSM50988, GSM50989, GSM50990, GSM50991, GSM50992, GSM50993, GSM50994, GSM50995, GSM50996, and GSM50997 [67].

Using the thermodynamically constrained Recon3D (Step 1) and the microarray datasets above, we executed the mCADRE routine. Briefly, as a first step, mCADRE scores those genes included in the Recon3D matrix according to their presence in the transcriptomic dataset. These scores are then attributed to reactions according to the initial model’s annotated gene-protein-reaction (GPR) relations. Next, the topology of the metabolic network is considered to update these scores. Once the procedure is complete, reactions with scores below a certain threshold are removed to complete the reconstruction of the cell-specific metabolic network. We refer our reader to [38] for a detailed explanation of the algorithm. Included in the supplementary files, we provide an updated file to be used in conjunction with mCADRE that admits Recon3D and works with a legacy version of IBM’s CPLEXstudio that supports a MATLAB bridge [68]. These files are necessary as the mCADRE algorithm requires CPLEX’s fast flux variability analysis - fastFVA, optimisation routine [69].

### Model reduction (Step 3)

In the previous steps, we have outlined how our GEMs, the BMMSC and Myeloma, were reconstructed to study the biochemistry occurring in human cells. In essence, these models can be used to probe the biochemistry of their respective metabolism. However, the validation of this approach’s results is limited because most experimental procedures focus on a small subset of cell metabolism. As a result, the model and its solutions leverage unnecessary complexity in its analysis, hindering a consistent and concise physiological representation. In our study, we are interested in the metabolism of cancer cells, and the data we can produce is centred around central carbon metabolism; this means we must keep those pathways that provide the energy, redox potential, and biomass precursors for cell growth and sustenance. We also include those pathways reported to be altered in cancer cells. Consequently, we specify the core subsystems of our models to be glycolysis, the pentose shunt, tricarboxylic acid cycle, oxidative phosphorylation, glutamate/glutamine metabolism, serine metabolism, urea cycle, and reactive oxygen species detoxification. We executed the redHuman algorithm, which performed a series of reductions [40]. Briefly, the core metabolic pathways defined above were specified in the routine. We then used the mCADRE GEM as input, which redHuman uses its subroutines, redGEM-lumpGEM, to reconstruct a reduced model around the core we specified [38, 40, 70, 71].

### Co-culture assembly, objective function, medium and phenotype specification (Step 4)

#### Co-Culture assembly and objective function

To model the experimental cell culture conditions, we must define an *in-silico* culture medium based on those used in the *in-vitro* counterpart. A correct definition of the medium in the model is fundamental for adequately representing intracellular metabolism. Using a combination of the redGEMX (part of redHuman) and the metabotools MATLAB package routine setMediumConstraints, we could guarantee that the reduced model has all the feasible pathways that consume and produce the components of the extracellular environment of the medium [40, 70, 72]. The redGEMX subroutine finds those pathways from the model needed to connect the extracellular metabolites to the core network defined above. We also employed constraints on the exchange fluxes to the extracellular medium in both cells by using data related to RPMI media composition. The advantage of using these methods in conjunction is that we were able to specifically tend to the needs of the reduced model whilst providing a physiologically sound *in-vitro* medium model that defines the medium components and those that can be uptaken by the cells but are not captured by the measured data such as 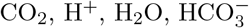, and NH_4_.

When considering the assembly of the *in-silico* co-culture model, we manually curated each cell-specific stochiometric matrix to reflect the compartments of each as unique (i.e., metabolite[cm] for myeloma cytosol and metabolite[bm] for BMMSC cytosol), excluding the extracellular fluxes which remain the same given that both cells share this space (specified in the model under the notation [e]). The consequent model spans a large metabolic matrix where duplicated extracellular sinks and sources were removed. Furthermore, the final model requires that both cells are equipped with biomass-producing reactions. We adhere to those specified by the Recon3D GEM [39]. There are several reasons, including experimental capabilities, for using this function as the optimisation objective of the model. One such argument was brought forward by a study which analysed and compared several CBM biomass functions for cancer metabolic modelling [4, 73]. Here, it was found that the formulation from the Recon family models (Recon 2 and Recon3D) leads to the most accurate results in terms of essential gene and growth rate predictions amongst all cancer cell lines they tested, including those for leukaemia research.

#### Multi-objective function

Our two cells residing within the microenvironment (i.e., the extracellular media) have different functions, each needing a different objective in a simulation. In a multi-objective paradigm, we assume that both cells, even under controlled environments, address various biological tasks (i.e., objectives) to sustain life. To obtain an optimal solution, we use the concept of “Pareto-optimality”. Briefly, a Pareto-optimal solution is one where the performance of one task would diminish the ability to achieve one or more other tasks. As we assume, for simplicity’s sake, that regardless of the biological functions of the two cell types, in our simulation, the biomass function carries sufficient knowledge about the biological system [74–76]. Our simulations implemented multi-objective optimisation using the MOFA routine for the COBRA package in MATLAB [77]. Briefly, MOFA maps the n-1-dimensional surface of the Pareto front, where n is the number of objectives, by calculating a large set of Pareto solutions. That is, the routine maps an area of multiple criteria decision-making in the presence of trade-offs between two or more conflicting objectives [77, 78].

#### Phenotype specification

Because metabolism is highly cell type- and state-specific, it offers valuable opportunities for diagnosing and treating cancers [79]. Mass spectrometry (MS) or nucleic magnetic resonance (NMR) based methods such as metabolomics measure relative abundance of cellular metabolites, which can help detect differences in metabolic state, but such abundance data does not inform on itself metabolic activities [41, 80, 81]. For a detailed assessment of intracellular metabolic activity in cells, stable isotope labelled nutrients are required [80–82]. This is because metabolic reaction rates, or fluxes, contribute to metabolic phenotypes and mechanisms of cellular regulation [41, 80, 82]. However, in most cases, isotopic labelling data cannot be interpreted intuitively due to the highly complex nature of atom rearrangements in metabolic pathways; instead, a formal model-based analysis approach is required to extract flux information from the labelling data. To quantify and characterise the metabolic phenotype in cells ^13^C-metabolic flux analysis (^13^C-MFA) is the gold standard for converting isotopic labelling data into corresponding metabolic flux maps [41, 42]. At a high-level ^13^C-MFA is formulated as a least-squares parameter estimation problem, where fluxes are unknown parameters that must be estimated by minimising the difference between the measured labelling data and labelling patterns simulated by the model, subject to stoichiometric constraints resulting from mass balances for intracellular metabolites and metabolite labelling states; the so-called mass isotopomers [41, 42].

Our main objective is to generate a quantitative map of cellular metabolism by assigning flux values to the reactions in an annotated metabolic network model and confidence intervals for each estimated flux. The confidence intervals act as lower and upper bounds in the CBM generated from steps 1 through 4. The result is the incorporation of experimental data and further modelling constraints. Consequently, we allow each cell in the model to acquire the phenotype displayed, or extrapolated, from the co-culture *in-vitro* setup. Although many routines can perform ^13^C-MFA, we have chosen the MATLAB package for isotopomer network compartmental analysis (INCA) [42]. Briefly, INCA uses the elementary metabolite unit (EMU) framework to simulate isotopic labelling in any arbitrary biochemical model. The routine is equipped with a Montecarlo parameter estimation routine, which we used to obtain confidence intervals for each predicted flux [41, 42, 83].

The selection of isotopic tracers in this study was ^13^C_6_-Glucose and ^13^C_5_-Glutamine. ^13^C_6_-Glucose is best for determining those fluxes in the upper sections of carbon metabolism, that is, glycolysis and the incorporation of its products into the TCA cycle [41]. ^13^C_5_-Glutamine, on the other hand, is used to determine exclusively lower parts, that is, the TCA cycle and reductive carboxylation. All tracing experiments performed were done in parallel, meaning that the cultures and initial conditions were identical, except that tracing was done in different batches. The data obtained from these experiments can also be incorporated in parallel because these isotopes are complementary [41, 42].

## Results

### Models in mono-culture: results and experimental validation

We performed benchmark tests to validate the predictive behaviour of both models, the BMMS cell and the MM cell. The following simulations were performed in mono-culture. Each cell-specific model was tested individually, where the extracellular medium was set to RPMI. We then performed, *in-silico*, experiments that address the consistency of cellular cultures [84, 85].

### Cell growth

Simulation for cell growth prediction rate was obtained by performing the Flux Balance Analysis routine with maximisation of biomass as the optimisation objective [47]. BMMSC growth rate has been reported to largely depend on cell culture conditions, including medium composition and its changes over time. Our experimental growth rate data from the HS-5 cell line cultured in RPMI medium was calculated to be 0.024 ±0.0097 /hour. Several works of literature report the *in-vitro* BMMSc population growth rate to be 0.027 /hour. Our bone marrow mesenchymal stem cell model predicts a growth rate of 0.0306 /hour. This was found by noting that each gram of dry cell weight (grDW) is assumed to be equivalent to one millimole of biomass. This means that the optimal *in-silico* specific growth rate is equal to [64]:

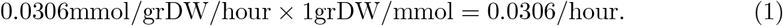

Our result is within the measured error of our experiments and those reported in literature (Fig. 3A) [64, 86–88]. We repeated the procedure to calculate the growth rate of the myeloma cell (Eq. 1). The model predicts the *in-silico* myeloma cell’s growth rate is 0.0265 /hour which agrees with our experimental results which indicate a growth of 0.02843 /hour (Fig. 3B) and those reported in literature [89–91].

**Fig 3.**
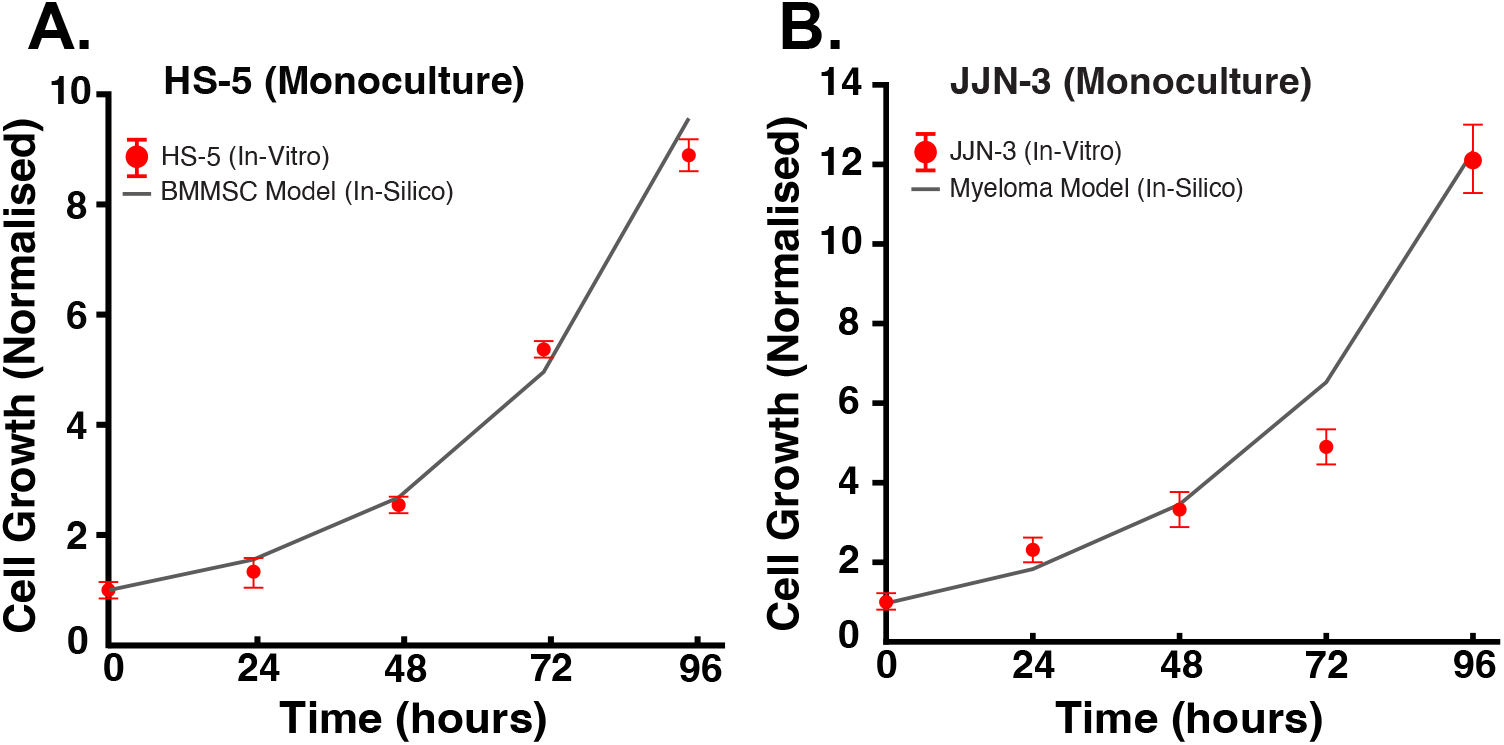
Growth rate comparison between model and experimental results in mono-culture. **A**. The measured normalised *in-vitro* culture growth of the HS-5 - a BMMSC cell line, in red, compared to the growth curve given by the exponential growth model solution fitted with the growth rate predicted by solving the BMMSC CBM using the FBA routine and maximum biomass production rate as the optimisation objective (in grey). **B**. The measured normalised *in-vitro* culture growth of the JJN-3 - a MM cell line, in red, compared to the growth curve given by the exponential growth model solution fitted with the growth rate predicted by solving the BMMSC CBM using the FBA routine and maximum biomass production rate as the optimisation objective (in grey).

### Oroboros respiration study

Because carbon metabolism is central to cancer study, as described in the methodology section, we tested each model’s (i.e., mono-culture) respiration by simulating an *in-silico* Oroboros study [32]. We then compared those performed in the *in-vitro* culture (also in mono-culture). Briefly, an Oroboros respiration study consists of a series of electron transport chain perturbations via pharmacological inhibitors to measure the respiratory capacity of the cell [32]. The procedure commences after basal measurements are recorded. This was achieved by measuring the oxygen (O_2_) flux when performing an FBA optimisation routine, where the objective function (Biomass) was set to find the minimum flux distribution through the network (Fig. 4 experiment-A. & C. *α*; model-& C *α*). The second step in the respiration study involves exposing cells to Oligomycin, an inhibitor of the ATP synthase (or complex V of the electron transport chain). By inhibiting the proton (H+) flux through this enzyme, the effects of Oligomycin cause an increased proton gradient across the mitochondrial inner membrane preventing electron transport through complexes I-IV (Fig. 4 experiment-A. & C. *β*; model-B. & C *β*). Oxygen consumption falls proportionally. The remaining rate of mitochondrial respiration represents a proton leak; that is, the protons pumped during electron transport that result in oxygen consumption but not ATP production (Fig. 4 experiment-A. & C. *δ*; model-B. & C *δ*). An increase in ATP-linked respiration, a measure of the cell’s capacity to meet its energy demands, would indicate an increase in ATP demand. In contrast, a decrease would indicate either low ATP demand, a lack of substrate availability, including severe damage to the oxidative phosphorylation pathway, which would impede the flow of electrons and result in a lower oxygen consumption rate (Fig. 4 experiment-A. & C. *ϵ*; model-B. & C *ϵ*). In both models, we achieved this by constraining the ATP synthase flux to zero and constraining the electron transport chain complexes to prevent reversal. Then we performed a second FBA optimisation routine, where the objective function was to maximise the biomass production rate.

**Fig 4.**
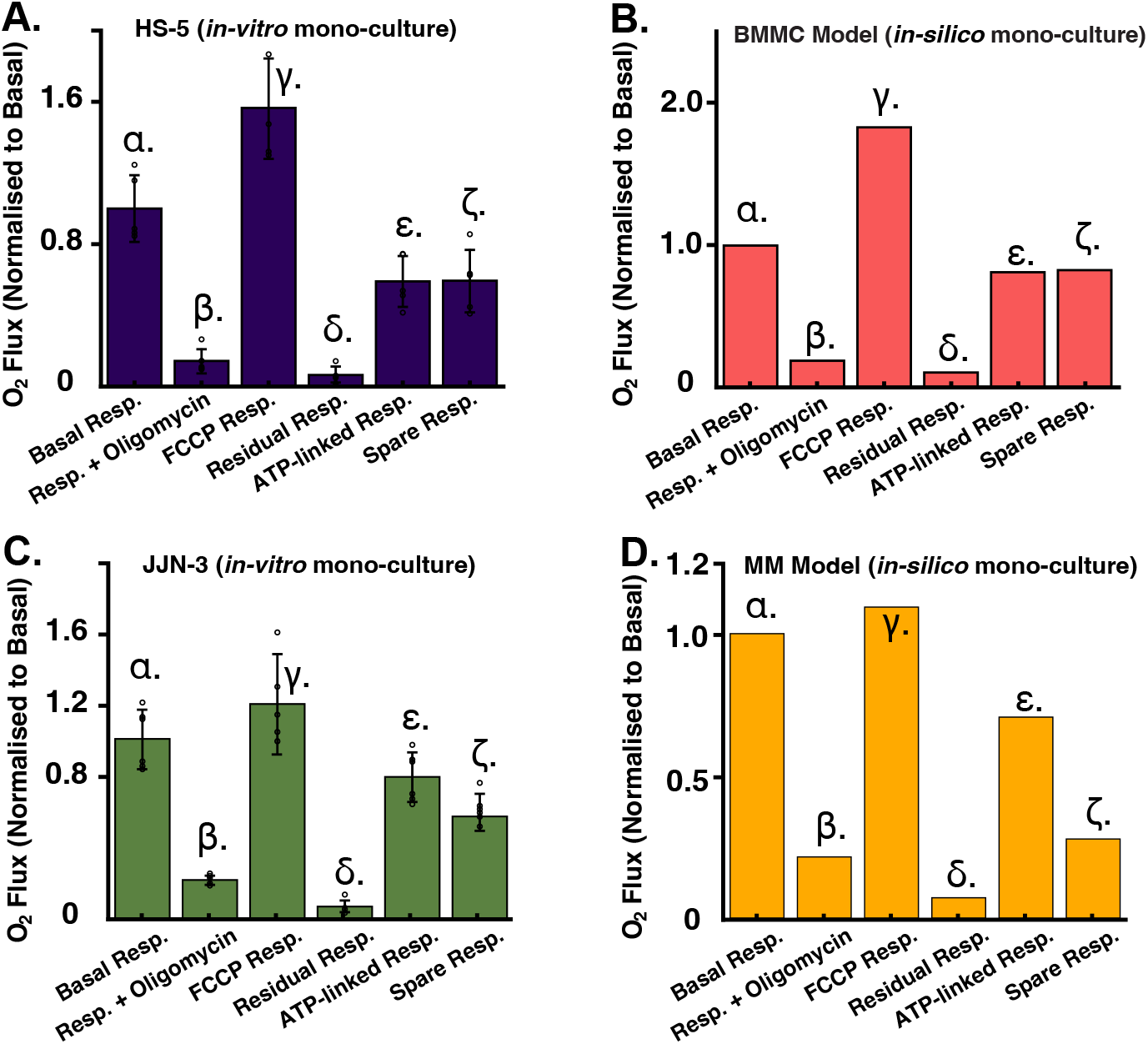
Mitochondrial respiration using the Oroboros respirometer is widely used to estimate mitochondrial respiration capacity. **A. & B**. Depict experimental results of HS-5 and JJN-3, BMMSC and MM cell-lines, respectively, mitochondrial respiration variability when perturbed with a series of pharmacological agents. **C. & D**. depict the experimental results of performing the *in-silico* version of the pharmacological perturbations in both cell models. These were achieved by constraining respective fluxes associated with the electron transport chain. For instance, Oligomycin is a potent ATP synthase inhibitor. To reproduce its effects on the model, we set the upper and lower bounds of the ATPsynthatse associated flux to zero, hence blocking any flux through that particular edge of the metabolic network. We then tracked the changes in mitochondrial O_2_ flux.

The following experiment involves FCCP, a potent uncoupling agent that collapses the proton gradient and disrupts the mitochondrial membrane potential. This agent is added following the cell’s exposure to Oligomycin. As a result, electron flow through the electron transport chain is uninhibited, and oxygen consumption by complex IV reaches the maximum. A high FCCP-stimulated oxygen consumption rate compared to a basal rate indicates that the mitochondria are using less than the maximal rate of electron transport supported by substrate supply from the cells. We performed this simulation by relaxing the constraints in the electron transport chain-related fluxes. We then performed a third FBA optimisation routine, where the objective was to maximise the flux through complex IV of the model’s electron transport chain (Fig. 4 experiment-A. & C. *γ*; model-B. & C *γ*). We then used the FCCP-stimulated oxygen consumption rate to calculate the cell’s spare respiratory capacity. This rate is defined as the difference between maximal respiration and basal respiration. Spare respiratory capacity measures the cell’s ability to respond to increased energy demand or under stress.

The last experiment involves exposing cells to a mixture of Rotenone, a complex I inhibitor, followed by the addition of antimycin A, an electron transport chain complex III inhibitor. This combination shuts down mitochondrial respiration and enables the calculation of nonmitochondrial respiration driven by processes outside the mitochondria. We performed this *in-silico* experiment by constraining the flux through complexes 1 and 3 of the electron transport chain. We performed an FBA optimization routine where biomass was used as the objective function to maximise (Fig. 4 experiment-A. & C. *ζ*; model-B. & C *ζ*).

### Models in co-culture: results and experimental validation

#### Co-culture cell growth

Following the experimental validation of the mono-culture models, we verified the co-culture *in-silico* model growth rate prediction. We found its value to be 0.0335 /hour for both cells, the BMMSCs and MM cells alike (Fig. 5). Interestingly, although it appears remarkably close to the measured values in its *in-vitro* counterpart, we cannot accurately validate the growth rate for the JJN-3 cell line in co-culture. This is because our experimental data was only performed for one biological replicate, rendering its statistical significance null. Because the experimental data appears inconclusive, we assume the model’s prediction suffices as a reasonable approximate growth according to those found in mono-culture, and those seen in literature [64, 86–91].

**Fig 5.**
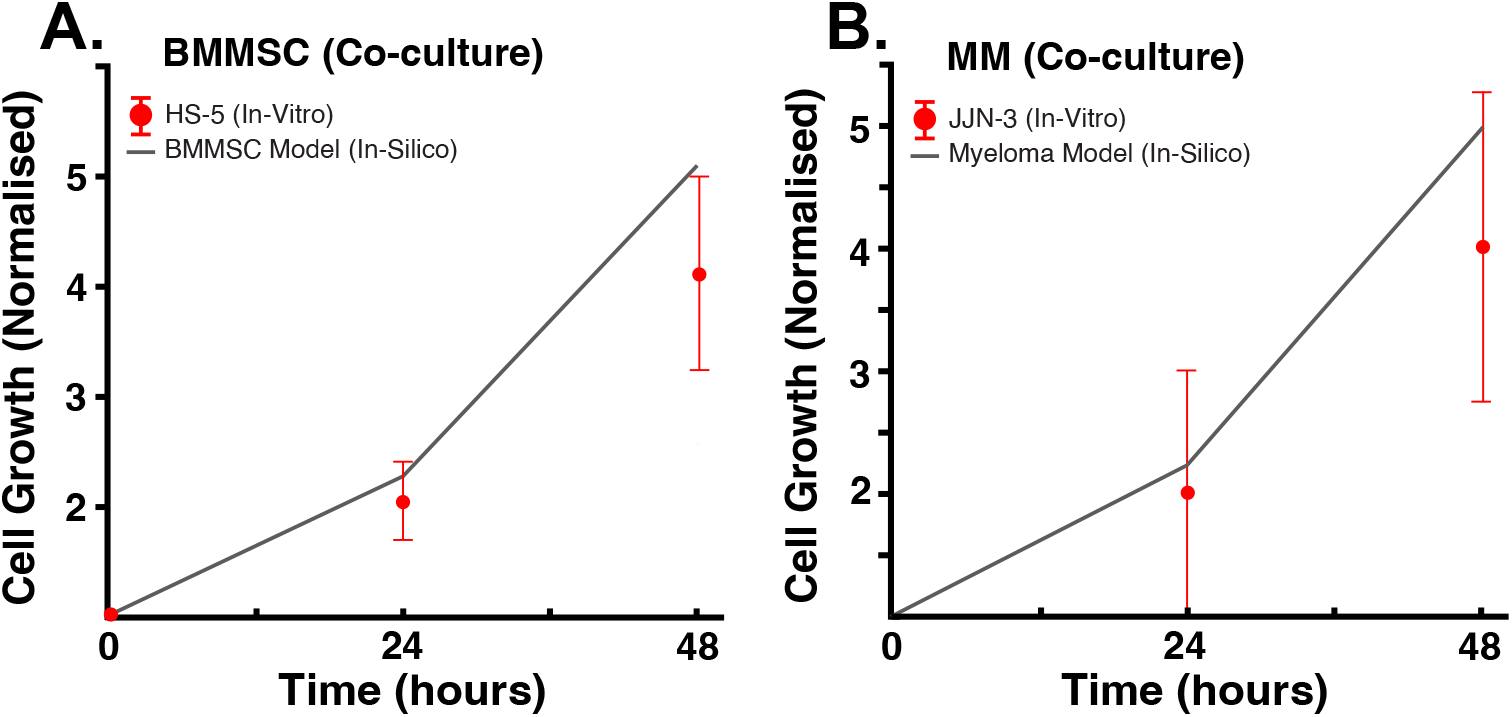
Growth rate comparison between model and experimental results in co-culture. **A**. The measured normalised *in-vitro* co-culture growth of the HS-5 - a BMMSC cell line, in red, compared to the growth curve given by the exponential growth model solution fitted with the growth rate predicted by the *in-silico* co-culture model (in grey). **B**. The measured normalised *in-vitro* co-culture growth of the JJN-3 - a MM cell line, in red, compared to the growth curve given by the exponential growth model solution fitted with the growth rate predicted by the in-silico co-culture model (in grey).

#### ^13^C-MFA

Mass and isotopomer balances were simulated for Myeloma cell and Bone Marrow Stem Cell, HS-5 and JJN-3 cell lines. Isotopic steady state was verified by measuring mass isotopomer distributions (MIDs) equilibration at the time points analysed. The isotopic steady state is reached when the change in isotopic enrichment over time falls within the measurement uncertainty range. We implemented the ^13^C_6_Glucose and ^13^C_5_Glutamine datasets in parallel datasets by regressing simultaneously to yield one complete metabolic flux map for each medium formulation. Glucose uptake flux was calculated to be 0.45 *µ*mol/10^6^ Cells/hr, and the measured glutamine was 0.63 *µ*mol/10^6^ Cells/hr - these values were calculated according to the protocol outlined in [41]. All fluxes were simulated in the INCA routine for MATLAB using a minimum of 100 unique restarts from random initial values to ensure a global minimum was found [42]. Flux results were subjected to a chi-square statistical test to assess goodness-of-fit, and 95% confidence intervals were calculated for each estimated flux value [42]. Fig. 6 summarises the results of our metabolic flux analysis. These results are thoroughly discussed in the next section.

**Fig 6.**
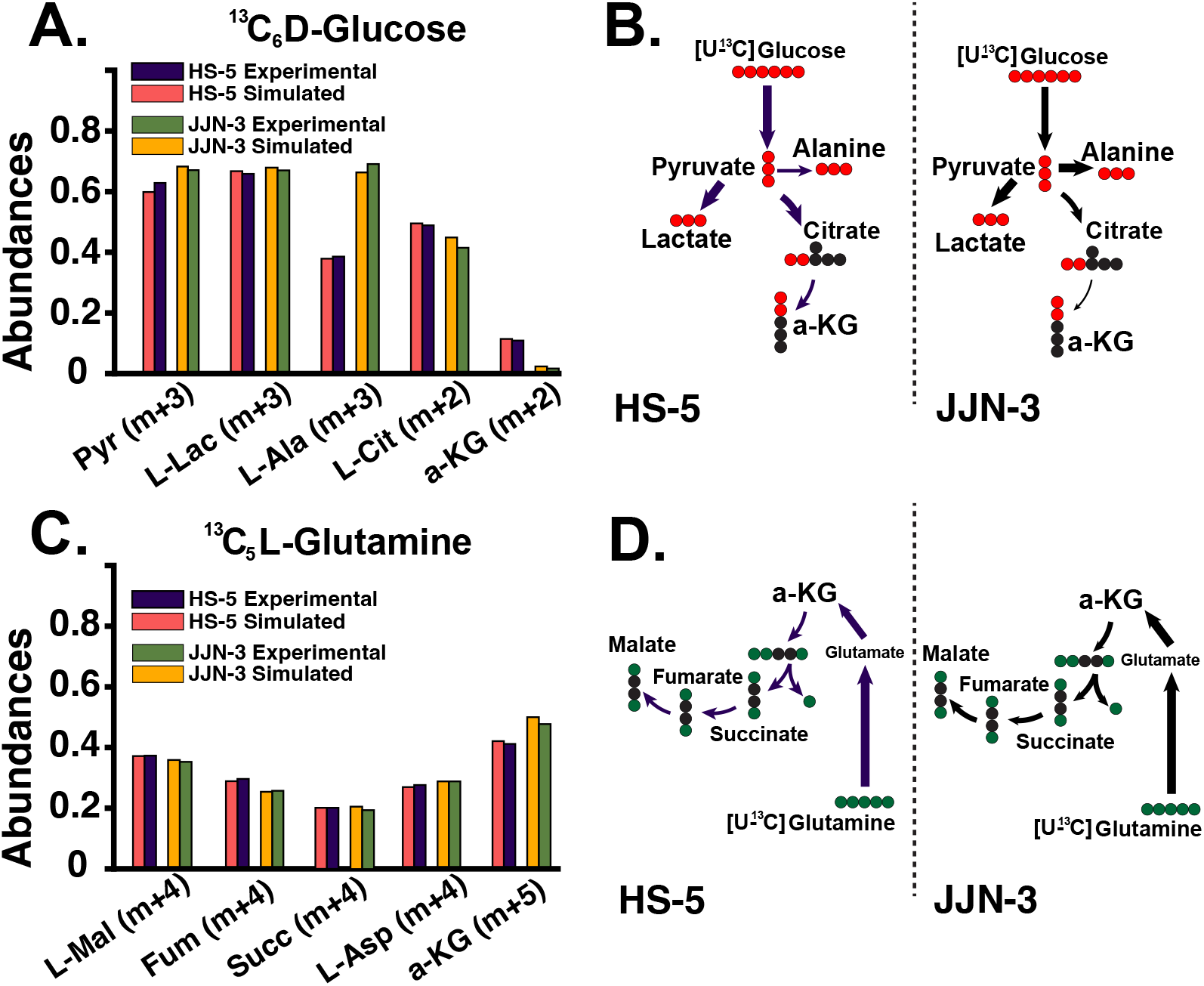
^13^C-MFA simulations. **A. & B**. Mass isotopomer distributions from ^13^C_6_Glucose and ^13^C_5_Glutamine tracing. Glucose-derived metabolite abundances. Data suggests high glycolytic activity in both cells. In JJN-3 cells, pyruvate-derived alanine is much higher than in HS-5 cells. The abundance of pyruvate in JJN-3 cells is also more significant than that of HS-5. It is possible to see that although pyruvate-derived citrate follows the expected TCA cycle, it does not continue downstream as a-ketoglutarate is low in JJN-3 cells. This indicates an interruption of the TCA cycle. **C. & D**. Mass isotopomer distributions from [U-13C]Glucose and [U-13C]Glutamine tracing. Glutamine-derived metabolite abundances. Data suggests that the TCA cycle in both cells is supported via anaplerosis, where exogenous glutamine is interconverted to a-ketoglutarate and then incorporated into the TCA cycle.

Please refer to the supplementary material file for the model used in our ^13^C-MFA simulations.

#### Co-culture model predictions

Once the co-culture model was curated, we performed flux balance analysis using the average between minimising and maximising the objective function. In doing so, we could generate a robust flux distribution to yield quality predictions while maintaining the critical functionality of those pathways required for biomass synthesis. Our *in-silico* simulation results reveal that both cells decrease their glycolytic activity as they go from mono-culture to co-culture. This, in principle, is a counterintuitive result as a high glycolytic activity is a hallmark of cancer metabolism. Therefore, we expected to see an enhancement of this behaviour when assembled in a more physiologically relevant way. However, this result capitulates the observed heterogeneity between cell types in mono-culture, as opposed to its results in co-culture. For instance, the mono-culture solutions of our models prioritise those fluxes associated with the glycolysis pathways to generate most of the ATP required for biomass synthesis. In contrast, we see this to a lesser extent in *in-vitro* co-cultured cell-lines. Contrary to our expectations, our mono-culture models support this experimental evidence where the in-vitro co-culture cell lines consume less glucose than mono-cultured. However, glucose uptake simulations performed in our co-culture model do not re-capitulate this behaviour, perhaps explaining the slightly faster cell growth rate result observed in Fig. 5. The glucose usage phenotype that the *in-vitro* co-culture exhibits was acquired once the ^13^C-isotope labelling data was incorporated.

It has been suspected that synthesising large amounts of bioenergetic metabolites requires malignant cells to establish a cooperative intercellular metabolic network. Our model predicts that a critical component of this mechanism is driven by the unidirectional export of pyruvate and lactate from bone marrow mesenchymal stem cell to the myeloma cell. Our simulations also indicate that this now exogenous pyruvate is uptaken by the myeloma cell to support its respiratory and biomass needs. In Fig. 8, it is depicted how in co-culture, BMMSC produces relatively large amounts of pyruvate relative to those it uses for its functionality. Indeed, a fraction of this branching point metabolite is destined for cell respiration and biomass (i.e., TCA cycle, alanine synthesis), whilst the rest is exported to the co-culture medium. Soon after pyruvate is incorporated into the myeloma cell’s metabolic network, it is destined to support macromolecule synthesis, which we suspect is mainly destined for fatty acids.

Indeed, we can show that in-vitro co-cultured BMMSCs and myeloma cells consume less glucose than when in mono-culture (Fig. 7 C.). These observations suggest that these two cell types work as a metabolic community that promotes more efficient nutrient use within the bone marrow (Fig. 7 A.). Experiments also revealed that co-cultured HS-5 cells showed increased glucose-derived pyruvate, while little change was observed in myeloma cells (Fig. 7 B.). This indicates that pyruvate may be trafficked between cell types, resulting in the enhanced efficiency of glucose metabolism, hence validating the model’s prediction.

**Fig 7.**
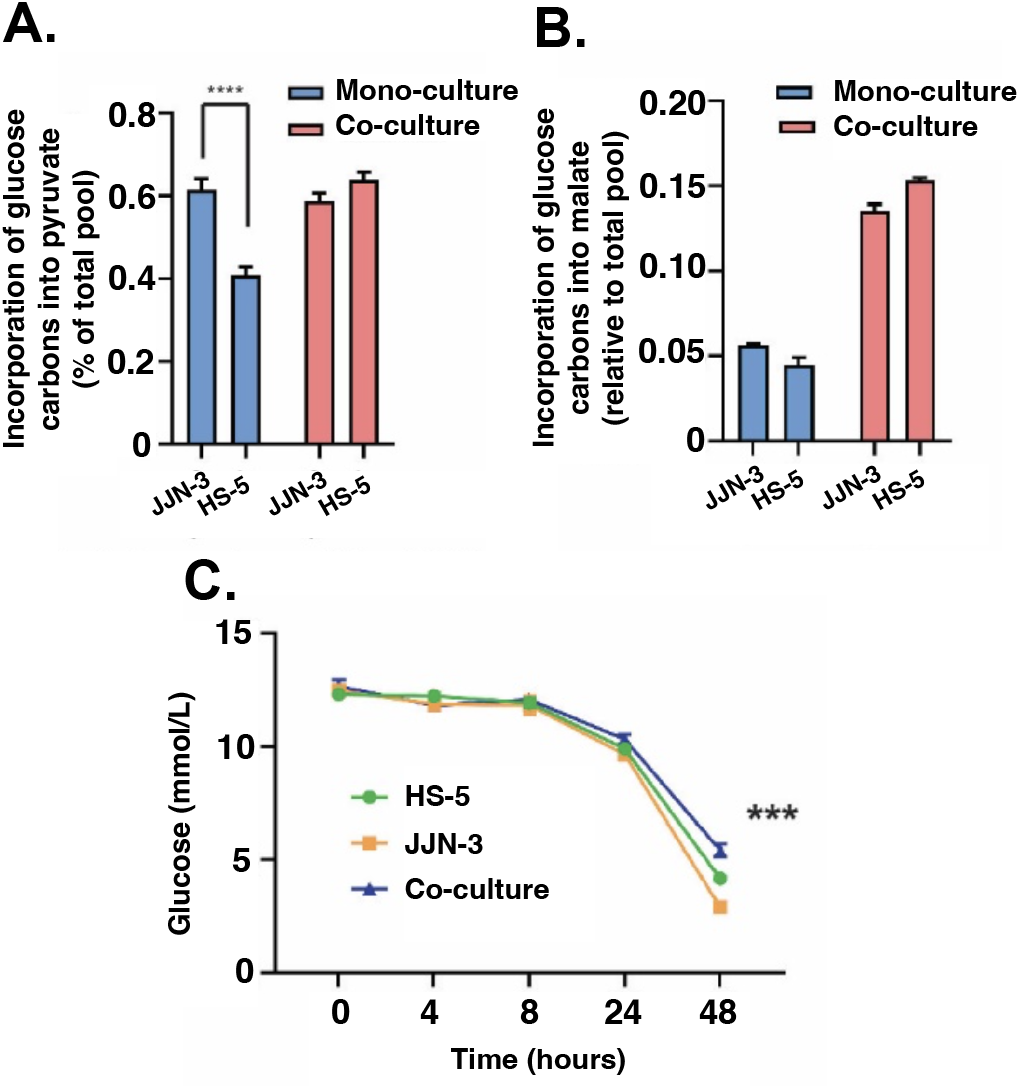
**A**. Incorporation of glucose carbons into pyruvate is also altered by co-culture; while HS-5 cells show lower pyruvate incorporation in mono-culture compared to co-culture conditions, there is little change in JJN-3 incorporation. **B**. When incubated with ^13^C_6_Glucose, label enrichment into the TCA cycle metabolite malate in both the myeloma cell line (JJN-3) or BMMSCs (HS-5) indicates a significant increase in oxidative glucose metabolism (m+2 isotopomer) as well as anaplerotic use (m+3 isotopomer). **C**. Co-culture of BMMSCs and MM cell lines, HS-5 and JJN-3, respectively, results in reduced glucose use compared to either cell type in mono-culture (corrected to cell number), suggesting a more efficient use of glucose carbons when both cells are cultured together. (In-figure referencing: ***; p*<*0.001, ****;p*<*0.0001. From Anova and Dunn’s multiple comparison).

**Fig 8.**
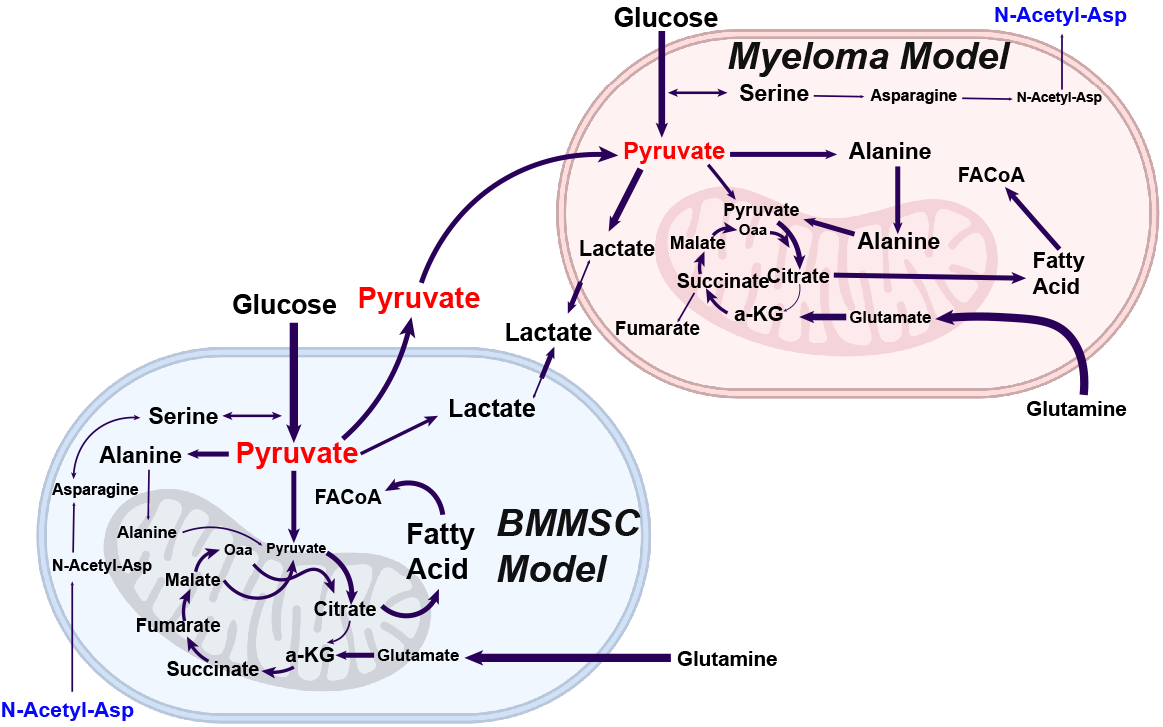
Model predictions. In co-culture, BMMSC produces relatively large amounts of pyruvate, with a fraction destined for cell respiration and the rest exported to the co-culture medium. Soon after, this pyruvate is uptaken by myeloma cells and incorporated into its carbon metabolism axis, destined to support biomass synthesis. However, both cells display a fractionated TCA cycle, in which L-citrate is exported, primarily for fatty acid synthesis, and the cycle is rescued via glutaminolysis. The model exhibits significant acetylation of asparagine to form N-acetyl-l-asparagine in myeloma cells. The metabolite is exported and uptaken by bone marrow mesenchymal stem cells, where reverse acetylation produces asparagine and acetate. This is used by the bone marrow mesenchymal stem cell for exogenous serine exchange, which is pipelined, in part, to produce pyruvate, lipids, antioxidants, and biomass.

The model’s flux distribution f exhibits that the ratio of pyruvate over lactate in HS-5 cells is asymmetric towards pyruvate, in agreement with our co-culture experiments. It is unclear, however, what is the destiny of the lactate. This is because the export and import of this metabolite are balanced with its synthesis to pyruvate and back, presumably to maintain the cytosolic redox balance of NAD+/NADH.

Once pyruvate is inside the mitochondria, it is catabolised mainly via pyruvate dehydrogenases (PDH) and to a lesser extent via the pyruvate carboxylases (PC), where it follows the TCA cycle terminating as L-Citrate, catalysed by the citrate synthase (CS). At this point, the model indicates that the metabolite is extruded from the mitochondria via citrate transport proteins in exchange for *±*-ketoglutarate, which we suspect is destined to synthesise fatty acids. As this flux is relatively large, the model bypasses the conversion of citrate to isocitrate by the aconitate hydratases, resulting in a relatively low proportion of mitochondrial glucose-derived a-ketoglutarate. These observations agree with our experimental data, in which we saw a decrease in the m+2 isotopomer from citrate to a-ketoglutarate (Fig. 6 B), indicating that mitochondrial a-ketoglutarate in BMMSCs cells, in its majority, is not glucose-derived. The model predicts that this extrusion aids the balance of *α*-ketoglutarate fluxes from mitochondrial citrate exchange at a steady state -previous studies of inhibition of aconitate hydratases due to high levels of ROS saw similar results [92]. According to our simulations, the BMMSC’s TCA cycle, beyond the L-citrate step, is rescued via anaplerotic reactions due to high glutaminolytic activity, a hallmark of cancer (Fig. 6 D). Finally, in our *in-silico* model, we also observed a considerable flux by the L-lactate dehydrogenases (LDH) from glucose-derived pyruvate, indicating that the BMMSC is also highly glycolytic; as expected (The Warburg effect). This glucose-derived L-lactate is then extruded. However, we cannot rule out cells re-import this lactate once extruded. The result may support previous studies where these cells observed a paracrine-like behaviour, where L-lactate is reabsorbed.

Following pyruvate incorporation into the mitochondria by the myeloma cell, our *in-silico* model exhibits high rates of L-Citrate production, observed in our experimental results (Fig. 6). As in the BMMSC, pyruvate-derived L-Citrate is mainly exported to the cytosolic compartment in exchange for a-ketoglutarate (specifically via the mitochondrial carrier CTP-Slc25a1). Once in the cytosol, L-Citrate is mainly converted into fatty acids; the model’s flux distribution selects long-chain fatty acids (i.e., palmitate). These fatty acids are primarily destined for biomass production, while a relatively small proportion is recycled back into the mitochondria via *β*-oxidation and incorporated back into the TCA cycle (Fig. 8). As most glucose-derived L-Citrate is exported, this cell exhibits low fluxes of isocitrate present in the TCA cycle of the myeloma model. Remarkably, these are even lower than those observed in the BMMSC cell model. Within the myeloma cell GEM, most of the available pyruvate is destined for the mitochondria and transported passively by the mitochondrial membrane pyruvate carriers (MPCs). However, an intriguing result of the model’s flux distribution describes that a significant proportion is transaminated into L-alanine in the cytosol and then transported into the mitochondria via an unspecified mitochondrial transporter. Once inside the mitochondria, L-alanine is de-transaminated back into pyruvate. The result is intriguing, as L-alanine is also a biomass precursor. Similarly to the BMMSC, most of the *±* -ketoglutarate necessary to resume the TCA cycle is supplied via the conversion of mitochondrial glutamate to *±*-ketoglutarate by L-Glutamate dehydrogenases. Finally, unlike the BMMSC results, the myeloma cell’s flux distribution predicts that besides obtaining *±*-ketoglutarate from L-Citrate transport and glutaminolysis, the cell balances its mitochondrial *±*-ketoglutarate requirements by increasing the flux through the mitochondrial *±*-ketoglutarate/malate transporter (expressed by the Slc25a10/11 genes). This serves a double purpose, contributing further to sustaining anabolism and the redox balance needed to maintain the cell’s high glycolytic demand.

Finally, we investigated additional possible intercellular metabolic exchange nodes across the multicellular model. Our results exhibit significant acetylation of asparagine to form N-acetyl-l-asparagine in myeloma cells (Fig. 8). The metabolite is exported and uptaken by the BMMSC model, where reverse acetylation produces asparagine and acetate. In our model, this is used to exchange it for extracellular serine, which is destined, in part, to produce pyruvate, lipids, antioxidants, and biomass. As the myeloma cells show elevated levels of glutamine production for biomass synthesis and energy requirements, they might not be able to proliferate without exogenous asparagine. This indicates a demand which exceeds what can be generated through the GS/ASNS pathway. Asparagine, therefore, is N-acetylated using acetyl-CoA as the acetyl-donor to form N-acetyl-l-asparagine. In this way, the cell uses N-acetyl-l-asparagine derived-asparagine as a factor to regulate biomass synthesis and, perhaps, cell proliferation. The presence of this behaviour in our model is remarkable, as asparagine has been theorised to be a critical metabolite for proliferating cancer cells.

## Discussion

Tumour metabolism remains an exciting space in which to identify novel drug targets. However, studies of cancer cell metabolism alone run the significant risk of identifying redundant metabolic enzymes in the cancer cell due to a more comprehensive metabolic network within tumours, including the neighbouring cells [93]. Understanding the physiology of these metabolic features promises the creation of a targeted approach wherby intercellular metabolic cross-talk also represents an exciting space for therapeutic manipulation, restricting the tumoural metabolic network and decreasing the resilience of the cancer cell.

Although reprogrammed metabolic networks are not fully understood due to a myriad of experimental limitations, we can provide insights into the cellular metabolic status of cells through *in-silico* models [94–96]. The ability of data integration, along with cell specificity in these models, may provide a broader picture, proving an invaluable tool to experimentalists. Furthermore, our proposed pipeline (Fig. 2) is also relevant in studies where modelling cell metabolism within the context of the tumour microenvironment, which constitutes a complex adaptive ecological system, is critical, and due to the difficulty in experimentally probing the interactions between multiple cell types remains challenging [96]. Such methodology (Fig. 2) presents an alternative to known algorithms that sacrifice data integration for reductionist techniques.

We demonstrated the applicability of our proposed pipeline by addressing a known gap in the field of multiple myeloma. We reconstructed a state-of-the-art co-culture *in-silico* model to probe the tumour metabolism at the genome-scale whilst providing the necessary granularity to underpin the intercellular metabolic cross-talk between cancer and stroma in the context of the framework for constraint-based modelling. Our simulations saw our models’ maximal *in-silico* growth rates, exhibited in both mono-culture and co-culture, agree with those observed in their respective experimental counterparts and those observed in literature [64]. We carried out these simulations under the assumption that the biological objective of the cancer cells is typically to grow (Figs. 3 & 5). However, even though our model reproduces the experimentally measured cell growth, it is possible that these rates could vary depending on the culture medium or *in-vivo* conditions, and also considering that assuming maximal growth is quite a reductionist approach and a cell population may be responding to different environmental cues or stresses, altering the cell’s objectives [14]. For instance, in certain conditions, cells may aim to minimise glucose uptake and maximise ATP, a behaviour observed in experimental co-cultures [97]. This fact may explain the variety of Pareto-optimal solutions given by performing the MOFA routine [77]. We, however, presented in Fig. 5 those solutions in the Pareto front that agree with our experimental observations. At the same time, more experimental data with various distinct mediums must be tested to accurately assert the validity of the model’s predictions, as we recognise that there exists a lack of regulatory constraints in our model [14, 98]. To address this, we will explore the addition of proteomic data in future studies. Doing so has resulted in even better support for experimentally observed phenomena, mainly when modelling effects of perturbations in cell metabolism, such as drug metabolism [98].

Taking the model’s validations into account, we carried out what we refer to as “phenotype tuning”. This consists in integrating ^13^C labelled glucose and glutamine data to constrain the model and produce physiologically relevant constraints. We found that myeloma cells exhibit an effect characterised by the malignancy’s reliance on glycolysis, leading to significant amounts of lactate production - this would be consistent with many previous studies on multiple cancer cell types [99, 100]. For instance, it has been hypothesised that lactate is not only a by-product of a malignancy’s highly glycolytic metabolism but can be salvaged by other cell types for mitochondrial oxidative energy production [28, 101, 102]. Similarly, it has also previously been suggested that lactate could be supplied by nearby stromal cells for the same reason - the directionality depending on the relative redox state of each cell [28, 102]. Furthermore, our experimental results demonstrate and confirm that when in co-culture, BMMSCs and myeloma cells (HS-5 and JJN-3, respectively) form a metabolic community spanning across the tumour microenvironment where instead of lactate, as previously thought, pyruvate is trafficked from stroma to myeloma and not in the other direction (Figs. 7 and 8). This intercellular interaction may be arbitrated unequivocally by transporters with a higher affinity for pyruvate over lactate, thereby establishing that these mechanisms rely mainly on pyruvate and not lactate as previously hypothesised [28, 101]. Interestingly, as predicted by our computational model, the pyruvate imported by myeloma cells appears to support the cell’s anabolism and energy generation. If our *in-silico* model predictions are correct, the result suggests that an alternative mode of action may be at play, which is aimed at supporting cell viability under limited resource conditions, which leads to an optimised metabolic community. Furthermore, with pyruvate transport targeting, lactate kinetics could be a powerful modality to perturb and increase the myeloma cell’s vulnerability to conventional drug therapies. Such a target comes timely, as historically, it has been challenging to find a suitable therapeutic window when targeting multiple myeloma metabolism because most agents focus on phenotypes common to other normal tissues. This is evident from the years of unsuccessful trialling glycolytic inhibitors, and the side effects observed with the current anti-metabolite therapies such as 5-fluorouracil [103, 104]. These agents suffer from a lack of tumour specificity, often resulting in significant side effects [103, 104].

A renaissance of research into cancer metabolism over the past two decades has been catalysed by significant improvements in our ability to detect and quantify thousands of different metabolites, alongside a revolution in sequencing technology that led to the identification of tumour-driving mutations in metabolic enzymes [105, 106]. Our study is part of a renewed drive towards developing novel agents targeting tumour metabolism. In this study we proposed an alternative *in-silico* reconstruction pipeline and used it to generate the first testable, integrated *in-vitro*/*in-silico* model of the metabolic network formed by malignant plasma cells and BMMSCs. To our knowledge, the majority of mathematical models that have been developed based on *in-vitro* data were produced under experimentally controlled conditions, where the metabolism of the malignant plasma cell in MM has characterised cell lines growing in isolation. Despite this, we are more likely to define efficacious cancer-specific targets for novel therapy development if the experimental system better re-capitulates the environment of the BM. Our integrated multidisciplinary workflows that iteratively utilise bench-side research alongside mathematical modelling can be utilised to identify and test the most appropriate targets for future pharmaceutical intervention. Introduction of metabolomics, fluxomics and growth-related data along with other related omics data into the reconstruction and validation process of GEMs can help increase the accuracy and prediction power of any model generated using our pipeline [107]. Our workflow, as demonstrated, can be used to generate useful *in-silico* models that underpin difficult-to-see behaviours and predict their results alongside *in-vitro* experiments.

## Acknowledgements

The authors would like to acknowledge funding from Cancer Research UK to **D.A.T**., **E.V.-S**., and **C.E.-G**. (**C42109/A26982** and **C42109/A24747**). Figs. 1, 2 and 8 were created with BioRender.com under an academic license. Finally, we would like to acknowledge the support and resources of the Birmingham Metabolic Tracer Analysis Core.

## Contribution

**E.V.-S**. conceived the mathematical model, performed *in-silico* experimentation and analysis, and wrote the manuscript. **I.S.-G**., **C.E.-G**., **& K.L.E**. performed *in-vitro* experiments. **D.A.T**. conceived the project, organised, and managed the study’s *in-vitro & in-silico* experiments. **F.S**. provided technical advice concerning the *in-silico* modelling.

## Appendix

### 1 Flux balance analysis

A CBM is represented through a system of algebraic equations, where in the absence of constraints, the resultant network’s flux distribution may lie at any point in its solution space. To this end, physiologically sound metabolic fluxes are determined from several constraints, including reversibility, reaction stoichiometry, and gene-associated rules. The stoichiometry of a biochemical reaction is crucial, as it tells the relative number of moles on either side of a balanced reaction. In a CBM this is represented by considering the metabolic network of interest with *m* metabolites and *n* reactions, and drafting a stoichiometric matrix **S**, where entry *s*_*i,j*_ represents the stoichiometric coefficient of the *i*^*th*^ metabolite in the *j*^*th*^ reaction:

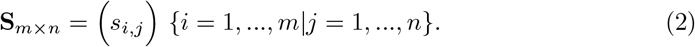

Using this formulation, we represent the steady-state of the system by defining a vector **v**, representing all reaction fluxes, satisfies

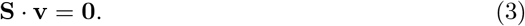

Note that the stoichiometric constraint of the model limits the system’s solution space, according to the mass balance concept.

Thermodynamical laws dictate the direction of every reaction from lower entropy to higher. Thus, a living organism encompassing the non-spontaneous reactions must have a continual energy input and then dissipate this energy in the form of unusable heat to guarantee continued existence. Imposing the constraints from this underlying concept constrains the solution space further, thus complying with thermodynamical laws. CBMs use flux limits for a given reaction, say, for example, the *i*^*th*^ reaction, to be always non-negative:

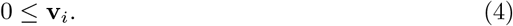

Mathematically, adhering to thermodynamical constraints rule out closed reaction cycles from model solution, and enforce the constraints on chemical potentials. Similarly, a lower (*lb*) or an upper (*ub*) flux bounds can be determined experimentally or theoretically for a specific reaction. In this case, constraints take the form of:

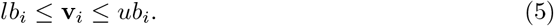

Just as constraints define the fluxes through a metabolic network, exact quantification of these fluxes cannot be achieved. However, assuming that metabolism has a particular predefined objective, a contested assumption of CBMs, it is possible to determine a flux distribution. This allows the system to satisfy an optimality criterion. Such criteria can vary depending on the application. A commonly assumed objective for flux balance models is the maximisation of biomass production. The approach allows determining ranges for the allowable flux distributions based on the hypothesised relevant biological objectives of the cell. Furthermore, proliferation occurs before differentiation. Thus in our study, biomass production flux is a choice for the objective function. The mathematical formulation of such an objective, accounting for defined flux restrictions with bounding limits, is given by:

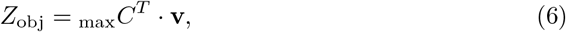

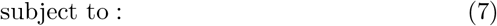

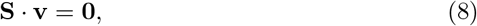

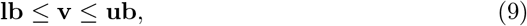

where *Z*_obj_ is the objective function and *C*^*T*^ is a vector of coefficients representing weights of fluxes in the objective [47].

### 2 *In-silico* culture medium

The medium composition as defined in our mono-cultures and co-culture models is given in Table 1, along with input parameters for the metabotools routine **setMediumConstraints**.

**Table 1.**
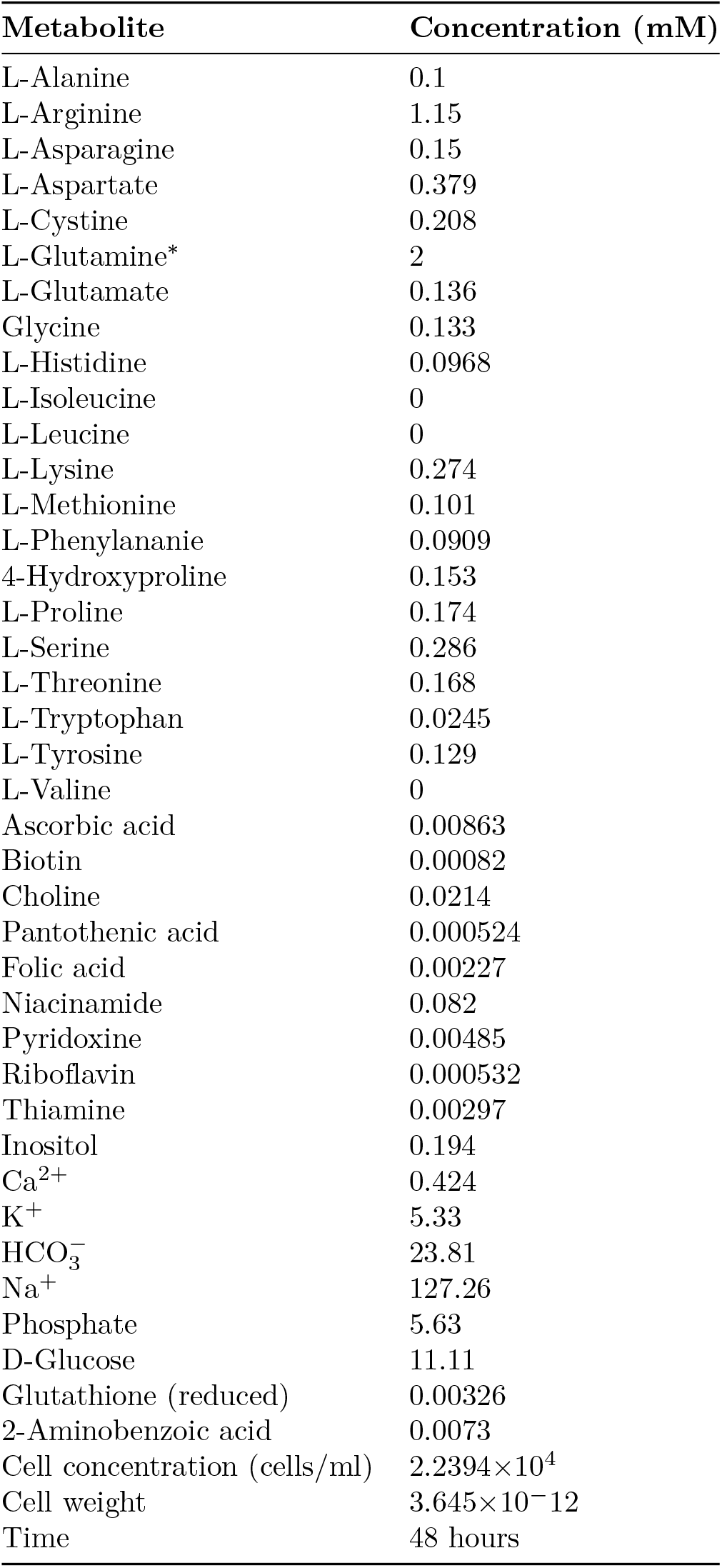
Medium Composition Parameters

| Metabolite | Concentration (mM) |
| --- | --- |
| L-Alanine | 0.1 |
| L-Arginine | 1.15 |
| L-Asparagine | 0.15 |
| L-Aspartate | 0.379 |
| L-Cystine | 0.208 |
| L-Glutamine* | 2 |
| L-Glutamate | 0.136 |
| Glycine | 0.133 |
| L-Histidine | 0.0968 |
| L-Isoleucine | 0 |
| L-Leucine | 0 |
| L-Lysine | 0.274 |
| L-Methionine | 0.101 |
| L-Phenylalanine | 0.0909 |
| 4-Hydroxyproline | 0.153 |
| L-Proline | 0.174 |
| L-Serine | 0.286 |
| L-Threonine | 0.168 |
| L-Tryptophan | 0.0245 |
| L-Tyrosine | 0.129 |
| L-Valine | 0 |
| Ascorbic acid | 0.00863 |
| Biotin | 0.00082 |
| Choline | 0.0214 |
| Pantothenic acid | 0.000524 |
| Folic acid | 0.00227 |
| Niacinamide | 0.082 |
| Pyridoxine | 0.00485 |
| Riboflavin | 0.000532 |
| Thiamine | 0.00297 |
| Inositol | 0.194 |
| Ca<sup>2+</sup> | 0.424 |
| K<sup>+</sup> | 5.33 |
| HCO<sub>3</sub><sup>-</sup> | 23.81 |
| Na<sup>+</sup> | 127.26 |
| Phosphate | 5.63 |
| D-Glucose | 11.11 |
| Glutathione (reduced) | 0.00326 |
| 2-Aminobenzoic acid | 0.0073 |
| Cell concentration (cells/ml) | 2.2394×10<sup>4</sup> |
| Cell weight | 3.645×10<sup>-12</sup> |
| Time | 48 hours |

