## Supplementary Material for "Mathematical reconstruction of the metabolic network in an *in-vitro* multiple myeloma model"

September 8, 2022

### 13C-MFA: Model

The $^{13}$C isotope labeling data generated by GC/MS analysis were analyzed using the MATLAB routine package INCA (publicly available at http://mfa.vueinnovations.com). Mass and isotopomer balances were simulated for Myeloma cell and Bone Marrow Stem Cell, HS-5 and JJN-3, cell lines. The simulations aim to infer fluxes of cell central carbon metabolism using an elementary metabolite unit (EMU) decomposition of the reaction network to simulate changes in isotope labeling induced by changes in metabolic fluxes. Isotopic steady state was verified by measuring equilibration of mass isotopomer distributions (MIDs) at the time points analyzed. Isotopic steady state is reached when the change in isotopic enrichment over time falls within range of the measurement uncertainty. We implemented the $^{13}C_6$Glucose and $^{13}C_5$Glutamine datasets in parallel datasets by regressing simultaneously to yield one complete metabolic flux map for each medium formulation. Fluxes were calculated using a minimum of 100 unique restarts from random initial values to ensure a global minimum was found. Flux results were subjected to a chi-square statistical test to assess goodness-of-fit, and 95% confidence intervals were calculated for each estimated flux value (included in INCA). The following is a summary of the model formulation, best-fit solutions, flux uncertainties, and goodness-of-fit metrics. The tracing was performed simulating the medium composition of the experimental proocedure where RPMI 1640 was used without branched-chain amino acids, L-leucine, L-isoleucine, and L-Valine. We then used the measured external D-glucose and L-Glutamine comncentrations to calculate their uptake fluxes as follows:

Assuming cells continuously grow, we use the exponential growth rate above. This was derived using:

$$N_x = N_{x,0} e^{\mu t}, \tag{1}$$

where $N_x$ is the number of cells, and $\mu$ is the growth rate. We then can obtain the doubling time by measuring the number of cells at two points. This is given by:

$$\mu = \frac{\ln\left(N_{x,t_2}\right) - \ln\left(N_{x,t_1}\right)}{\Delta t}, \tag{2}$$

the doubling time, then, is given by:

$$t_d = \frac{\ln(2)}{\mu}. \tag{3}$$

Table 1: $^{13}$C-MFA Model

| **Reaction** | **Formula + Atomic Transition** |
|---|---|
| R1 | glc.ext (abcdef) → glc (abcdef) |
| R2 | glc.ext (abcdef) → glc.m (abcdef) |
| R3 | gln.ext (abcde) → gln (abcde) |
| R4 | asp.ext (abcd) → asp (abcd) |
| R5 | val.ext (abcde) → val (abcde) |
| R6 | gly.ext (ab) → gly (ab) |
| R7 | ser.ext (abc) → ser (abc) |
| R8 | ala.ext (abc) ↔ ala (abc) |
| R9 | ala.med (abc) → ala.ext (abc) |
| R10 | o2.ext → o2 |
| R11 | o2.ext → o2.m |
| R12 | pro.ext (abcde) → pro (abcde) |
| R13 | lac.ext (abc) → lac (abc) |
| R14 | glc (abcdef) + atp → g6p (abcdef) + adp |
| R15 | g6p (abcdef) → f6p (abcdef) |
| R16 | f6p (abcdef) + atp → fbp (abcdef) + adp |
| R17 | fbp (abcdef) → dhap (cba) + gap (def) |
| R18 | dhap (abc) → gap (abc) |
| R19 | gap (abc) + adp + nad → 3pg (abc) + atp + nadh |
| R20 | 3pg (abc) → pep (abc) |
| R21 | pep (abc) + adp → pyr (abc) + atp |
| R22 | pyr (abc) + nadh ↔ lac (abc) + nad |
| R23 | lac (abc) → lac.ext (abc) |
| R24 | pyr (abc) → pyr.ext (abc) |
| R25 | pyr (abc) + nad.bm → accoa (bc) + co2 (a) + nadh.bm |

In this way, external metabolite rates were calculated as follows:

$$r_i = \frac{\mu V \Delta C_i}{\Delta N_x} \times 1000. \tag{4}$$

Here, $\Delta C_i$ (nmol/L) is the change in concentration of metabolite $i$, on the medium between the two time points, $\Delta N_x$ is the change in cell number, $V$ represents the volume of the cell culture, and $\mu$ is the growth rate. We multiply by 1000 to convert L to mL.

Table 2: $^{13}$C-MFA Model Continued...

| **Reaction** | **Formula + Atomic Transition** |
|---|---|
| R26 | pyr (abc) + co2 (d) + atp.bm → oaa (abcd) + adp.bm |
| R27 | accoa (ab) + oaa (cdef) → cit (fedbac) |
| R28 | cit (abcdef) + nad.bm ↔ akg (abcde) + co2 (f) + nadh.bm |
| R29 | 0.5*akg (abcde) + 0.5*akg (fghij) + nad.bm → ... |
| R29 | 0.5*succoa (bcde) + 0.5*succoa (jihg) + 0.5*co2 (a) + 0.5*co2 (f) + nadh.bm |
| R30 | 0.5*succoa (abcd) + 0.5*succoa (efgh) + adp.bm → ... |
| R30 | 0.5*succ (abcd) + 0.5*succ (efgh) + atp.bm |
| R31 | 0.5*succ (abcd) + 0.5*succ (efgh) + fad.bm → ... |
| R31 | 0.5*fum (abcd) + 0.5*fum (efgh) + fadh.bm |
| R32 | 0.5*fum (abcd) + 0.5*fum (efgh) → 0.5*mal (abcd) + 0.5*mal (efgh) |
| R33 | mal (abcd) + nad.bm → oaa (abcd) + nadh.bm |
| R34 | 0.5*o2 + nadh.bm + 3.5*adp.bm + 0.25*fadh.bm → ... |
| R34 | h2o + nad.bm + 3.5*atp.bm + 0.25*fad.bm |
| R35 | atp.bm + adp ↔ adp.bm + atp |
| R36 | gln (abcde) → glu (abcde) |
| R37 | h2o → |
| R38 | co2 (a) → |
| R39 | atp ↔ adp |
| R40 | pyr.ext (abc) → |
| R41 | lac.ext (abc) → |
| R42 | oaa (abcd) + glu (efghi) → asp (abcd) + akg (efghi) |
| R43 | asp (abcd) + glu.bc (efghi) ↔ asp.bc (abcd) + glu (efghi) |
| R44 | asp.bc (abcd) + akg.bc (efghi) ↔ oaa.bc (abcd) + glu.bc (efghi) |
| R45 | oaa.bc (abcd) + nadh ↔ mal.bc (abcd) + nad |
| R46 | mal.bc (abcd) + akg (efghi) ↔ mal (abcd) + akg.bc (efghi) |
| R47 | glu (abcde) + nad.bm ↔ akg (abcde) + nadh.bm |
| R48 | Fa.ext → Fa.bc |
| R49 | c1.ext → c1b |
| R50 | Facoa + 7*c2 (ab) + 6*nad.bm + 6*h2o + 6*fad.bm → ... |
| R50 | 7*accoa (ab) + 6*nadh.bm + 6*fadh.bm |

Table 3: $^{13}$C-MFA Model Continued...

| Reaction | Formula + Atomic Transition |
|---|---|
| R51 | cit (abcdef) → cit.bc (abcdef) |
| R52 | cit.bc (abcdef) + atp → accoa.bc (ab) + oaa (abcd) + adp |
| R53 | accoa.bc (ab) → Facoa.bc + co2 (a) + co2 (b) |
| R54 | g6p (abcdef) → ru5p (bcdef) + co2 (a) |
| R55 | ru5p (abcde) ↔ x5p (abcde) |
| R56 | ru5p (abcde) ↔ r5p (abcde) |
| R57 | x5p (abcde) ↔ ec2 (ab) + gap (cde) |
| R58 | f6p (abcdef) ↔ ec2 (ab) + e4p (cdef) |
| R59 | s7p (abcdefg) ↔ ec2 (ab) + r5p (cdefg) |
| R60 | f6p (abcdef) ↔ ec3 (abc) + gap (def) |
| R61 | s7p (abcdefg) ↔ ec3 (abc) + e4p (defg) |
| R62 | fum (abcd) → fum.ext (abcd) |
| R63 | fum.ext (abcd) → |
| R64 | val (abcde) ↔ akic (abcd) + co2 (e) |
| R65 | akic (abcd) ↔ succoa (abcd) |
| R66 | 3pg (abc) + nad + glu (defgh) → pser (abc) + nadh + akg (defgh) |
| R67 | pser (abc) → ser (abc) |
| R68 | ser (abc) ↔ gly (ab) + co2 (c) |
| R69 | gly (ab) → |
| R70 | pyr (abc) + glu (defgh) ↔ ala (abc) + akg (defgh) |
| R71 | ala (abc) → |
| R72 | mal (abcd) + nad.bm → pyr (abc) + co2 (d) + nadh.bm |
| R73 | oaa (abcd) → pep (abc) + co2 (d) |
| R74 | ala (abc) ↔ ala.bc (abc) |
| R75 | ala.bc (abc) + akg (defgh) ↔ pyr.bc (abc) + glu (defgh) |
| R76 | pyr.bc (abc) → pyr (abc) |
| R77 | pro (abcde) + fad.bm → 1pyr5c (abcde) + fadh.bm |
| R78 | 1pyr5c (abcde) → pro (abcde) |
| R79 | 1pyr5c (abcde) + nad.bm → glu (abcde) + nadh.bm |
| R80 | glu (abcde) + nadh.bm → 1pyr5c (abcde) + nad.bm |
| R81 | pro (abcde) → |
| R82 | glc.m (abcdef) + atp.m → g6p.m (abcdef) + adp.m |
| R83 | g6p.m (abcdef) → f6p.m (abcdef) |
| R84 | f6p.m (abcdef) + atp.m → fbp.m (abcdef) + adp.m |
| R85 | fbp.m (abcdef) → dhap.m (cba) + gap.m (def) |
| R86 | dhap.m (abc) → gap.m (abc) |
| R87 | gap.m (abc) + adp.m + nad.m → 3pg.m (abc) + atp.m + nadh.m |
| R88 | 3pg.m (abc) → pep.m (abc) |
| R89 | pep.m (abc) + adp.m → pyr.m (abc) + atp.m |
| R90 | pyr.m (abc) + nadh.m ↔ lac.m (abc) + nad.m |
| R91 | lac.m (abc) → lac.ext (abc) |
| R92 | pyr.ext (abc) ↔ pyr.m (abc) |
| R93 | pyr.m (abc) → |
| R94 | atp.m ↔ adp.m |
| R95 | pyr.m (abc) + nad.mm → accoa.m (bc) + co2.m (a) + nadh.mm |
| R96 | pyr.m (abc) + co2.m (d) + atp.mm → oaa.m (abcd) + adp.mm |
| R97 | accoa.m (ab) + oaa.m (cdef) → cit.m (fedbac) |
| R98 | cit.m (abcdef) + nad.mm ↔ akg.m (abcde) + co2.m (f) + nadh.mm |
| R99 | 0.5*akg.m (abcde) + 0.5*akg.m (fghij) + nad.mm →... |
| R99 | 0.5*succoa.m (bcde) + 0.5*succoa.m (jihg) + 0.5*co2.m (a) + 0.5*co2.m (f) + nadh.mm |

Table 4: $^{13}$C-MFA Model Continued...

| **Reaction** | **Formula + Atomic Transition** |
|---|---|
| R101 | 0.5*succoa.m (abcd) + 0.5*succoa.m (efgh) + adp.mm →... |
| R101 | 0.5*succ.m (abcd) + 0.5*succ.m (efgh) + atp.mm |
| R102 | 0.5*succ.m (abcd) + 0.5*succ.m (efgh) + fad.mm →... |
| R102 | 0.5*fum.m (abcd) + 0.5*fum.m (efgh) + fadh.mm |
| R103 | 0.5*fum.m (abcd) + 0.5*fum.m (efgh) →... |
| R103 | 0.5*mal.m (abcd) + 0.5*mal.m (efgh) |
| R104 | mal.m (abcd) + nad.mm → oaa.m (abcd) + nadh.mm |
| R105 | 0.5*o2.m + nadh.mm + 3.5*adp.mm + 0.25*fadh.mm →... |
| R105 | h2o.m + nad.mm + 3.5*atp.mm + 0.25*fad.mm |
| R106 | atp.mm + adp.m ↔ adp.mm + atp.m |
| R107 | h2o.m → |
| R108 | co2.m (a) → |
| R109 | gln.m (abcde) → glu.m (abcde) |
| R110 | oaa.m (abcd) + glu.m (efghi) → asp.m (abcd) + akg.m (efghi) |
| R111 | asp.m (abcd) + glu.mc (efghi) ↔ asp.mc (abcd) + glu.m (efghi) |
| R112 | asp.mc (abcd) + akg.mc (efghi) ↔ oaa.mc (abcd) + glu.mc (efghi) |
| R113 | oaa.mc (abcd) + nadh.m ↔ mal.mc (abcd) + nad.m |
| R114 | mal.mc (abcd) + akg.m (efghi) ↔ mal.m (abcd) + akg.mc (efghi) |
| R115 | glu.m (abcde) + nad.mm ↔ akg.m (abcde) + nadh.mm |
| R116 | Fa.ext → Fa.mc |
| R117 | c1.ext → c1.m |
| R118 | c2.ext (ab) → c2.m (ab) |
| R119 | Fa.mc + c1.m + atp.m → Facoa.mc + adp.m |
| R120 | Facoa.mc → Facoa.m |

Table 5: $^{13}$C-MFA Model Continued...

| Reaction | Formula + Atomic Transition |
|---|---|
| R121 | Facoa.m + 7*c2.m (ab) + 6*nad.mm + 6*h2o.m + 6*fad.mm → ... |
| R121 | 7*accoa.m (ab) + 6*nadh.mm + 6*fadh.mm |
| R122 | cit.m (abcdef) → cit.mc (abcdef) |
| R123 | cit.mc (abcdef) + atp.m → accoa.mc (ab) + oaa.m (abcd) + adp.m |
| R124 | accoa.mc (ab) → Facoa.mc + co2.m (a) + co2.m (b) |
| R125 | g6p.m (abcdef) → ru5p.m (bcdef) + co2.m (a) |
| R126 | ru5p.m (abcde) ↔ x5p.m (abcde) |
| R127 | ru5p.m (abcde) ↔ r5p.m (abcde) |
| R128 | x5p.m (abcde) ↔ ec2.m (ab) + gap.m (cde) |
| R129 | f6p.m (abcdef) ↔ ec2.m (ab) + e4p.m (cdef) |
| R130 | s7p.m (abcdefg) ↔ ec2.m (ab) + r5p.m (cdefg) |
| R131 | f6p.m (abcdef) ↔ ec3.m (abc) + gap.m (def) |
| R132 | s7p.m (abcdefg) ↔ ec3.m (abc) + e4p.m (defg) |
| R133 | fum.m (abcd) → fum.ext (abcd) |
| R134 | fum.ext (abcd) → |
| R135 | val.m (abcde) ↔ akic.m (abcd) + co2.m (e) |
| R136 | akic.m (abcd) ↔ succoa.m (abcd) |
| R137 | 3pg.m (abc) + nad.m + glu.m (defgh) →... |
| R137 | pser.m (abc) + nadh.m + akg.m (defgh) |
| R138 | pser.m (abc) → ser.m (abc) |
| R139 | ser.m (abc) ↔ gly.m (ab) + co2.m (c) |
| R140 | gly.m (ab) → |
| R141 | pyr.m (abc) + glu.m (defgh) ↔ ala.m (abc) + akg.m (defgh) |
| R142 | ala.m (abc) → |
| R143 | mal.m (abcd) + nad.mm → pyr.m (abc) + co2.m (d) + nadh.mm |
| R144 | oaa.m (abcd) → pep.m (abc) + co2.m (d) |
| R145 | ala.m (abc) ↔ ala.mc (abc) |
| R146 | ala.mc (abc) + akg.m (defgh) ↔ pyr.mc (abc) + glu.m (defgh) |
| R147 | pyr.mc (abc) → pyr.m (abc) |
| R148 | pro.m (abcde) + fad.mm → 1pyr5c.m (abcde) + fadh.mm |
| R149 | 1pyr5c.m (abcde) → pro.m (abcde) |
| R150 | 1pyr5c.m (abcde) + nad.mm → glu.m (abcde) + nadh.mm |
| R151 | glu.m (abcde) + nadh.mm → 1pyr5c.m (abcde) + nad.mm |
| R152 | pro.m (abcde) → |
